# Cryo-EM structures of the DCPIB-inhibited volume-regulated anion channel LRRC8A in lipid nanodiscs

**DOI:** 10.1101/442459

**Authors:** David M. Kern, SeCheol Oh, Richard K. Hite, Stephen G. Brohawn

## Abstract

Hypoosmotic conditions activate volume-regulated anion channels in vertebrate cells. These channels are formed by leucine-rich repeat-containing protein 8 (LRRC8) family members and contain LRRC8A in homo- or hetero-hexameric assemblies. Here we present single-particle cryo-electron microscopy structures of LRRC8A in complex with the inhibitor DCPIB reconstituted in lipid nanodiscs. DCPIB plugs the channel like a cork in a bottle - binding in the extracellular selectivity filter and sterically occluding ion conduction. Constricted and expanded structures reveal coupled dilation of cytoplasmic LRRs and the channel pore, suggesting a mechanism for channel gating by internal stimuli. Conformational and symmetry differences between LRRC8A structures determined in detergent micelles and lipid bilayers related to reorganization of intersubunit lipid binding sites demonstrate a critical role for the membrane in determining channel structure. These results provide insight into LRRC8 gating and inhibition and the role of lipids in the structure of an ionic-strength sensing ion channel.

## Introduction

Volume control is vital for a cell to respond to its environment. Without volume regulation, vertebrate cells are at the mercy of osmotic imbalance, a property utilized at extremes for biochemical cell lysis. However, by exporting or importing osmolytes in response to environmental or intracellular cues, cells can withstand osmotic stresses and actively regulate volume during cell growth, migration, and death.

Vertebrate cells respond to hypotonic environments by opening channels for chloride and other anions, permitting exchange of diverse osmolytes and requisite water across the membrane to alleviate osmotic imbalance (Jentsch, 2016). The underlying channels, termed volume-regulated anion channels (VRACs) (Nilius et al., 1997), have variable selectivity and conductance between cells, but are similarly activated by osmotic stimuli and blocked by small molecule anionic inhibitors including DCPIB (4-[(2-Butyl-6,7-dichloro-2-cyclopentyl-2,3-dihydro-1-oxo-1H-inden-5-yl)oxy]butanoic acid) (Decher et al., 2001). While apparently ubiquitous in vertebrate cells, these channels lacked a molecular identity until 2014, when leucine-rich repeat-containing family member 8A (LRRC8A, also named SWELL1) was identified as a required component of VRAC in cells (Qiu et al., 2014; Voss et al., 2014). LRRC8A and its paralogs LRRC8B, C, D, and E were proposed to form hexameric ion channels through their homology to Pannexins in the transmembrane helix-containing region (Abascal and Zardoya, 2012).

VRACs have been implicated in a wide array of physiological and pathophysiological processes including insulin signaling in adipocytes (Zhang et al., 2017), neurotransmitter release from astrocytes and brain damage after stroke (Hyzinski-García et al., 2014; Lutter et al., 2017), passage of the chemotherapeutic cisplatin into cancer cells (Planells-Cases et al., 2015), and osmotic control during spermatogenesis (Lück et al., 2018). A chromosomal translocation in humans that truncates LRRC8A in the LRR region is associated with the B-cell deficiency agammaglobulinemia (Sawada et al., 2003). A mouse knockout of *LRRC8A* exhibits increased mortality and developmental defects in addition to significant defects in T cell development and function (Kumar et al., 2014). The broad expression of LRRC8s in vertebrate cells suggests VRACs may have additional, yet-undiscovered, roles in cell biology and human physiology.

Functional expression of VRAC in cells requires LRRC8A (Qiu et al., 2014; Voss et al., 2014). LRRC8A can form homomeric channels as well as heteromeric channels with its LRRC8B, C, D and E paralogs; channels have been shown to contain one, two, or three different LRRC8 family members (Lutter et al., 2017). Channel properties, including ion selectivity and conductance, are modulated by subunit composition and the diversity of native VRAC properties is presumably a consequence of the different combinations of homomeric and heteromeric channels expressed in different cells. For example, LRRC8A homomeric channels exhibit low conductance relative to heteromeric channels (Deneka et al., 2018; Kasuya et al., 2018; Kefauver et al., 2018a; Syeda et al., 2016) and only LRRC8AD heteromers readily conduct larger molecules such as cisplatin and the antibiotic blasticidin S (Lee et al., 2014; Planells-Cases et al., 2015). Mechanistically, LRRC8 channels are opened by low cytoplasmic ionic strength, but the precise molecular mechanisms for sensing internal ionic strength and transmitting this stimulus to the opening of a channel gate are unknown (Syeda et al., 2016; Voets et al., 1999).

Here we report structures determined by cryo-electron microscopy of the homohexameric LRRC8A channel embedded in lipid nanodiscs and bound to the inhibitor DCPIB. The structures reveal the architecture of the LRRC8 family in a lipid environment, the mechanism of channel inhibition by DCPIB, and provide insight into the roles of bound lipids in stabilizing channel formation. LRRC8A channel complexes display significant structural heterogeneity, which we resolve into constricted and expanded states associated with conformationally heterogenous LRR domains. We propose these differences are related to ionic strength sensing and gating conformational changes. We further compare the structures in lipid nanodiscs to structures in the detergent digitonin presented here and in three recent reports (Deneka et al., 2018; Kasuya et al., 2018; Kefauver et al., 2018a). Differences in observed symmetry and LRR conformation between nanodisc-embedded and detergent-solubilized channels suggest an integral role for the membrane environment in LRRC8A structure.

## Results

### Structure of LRRC8A in lipid nanodiscs

LRRC8s are endogenously expressed in vertebrate cells typically used for protein overexpression, so we expressed and purified mouse LRRC8A from *Spodoptera frugiperda* (SF9) insect cells to ensure a homogenous channel preparation for further study (Figure 1-supplement 1). To assess the activity of channels from this preparation, we reconstituted LRRC8A purified in detergent into phosphatidylcholine lipids and recorded from proteoliposome patches. As is the case in cells (Syeda et al., 2016), channel activity was only observed in low ionic strength solutions (e.g. 70 mM KCl, Figure 1A). Channels displayed asymmetric conductance at positive and negative potentials with similar open probability. Purified LRRC8A therefore retains the characteristic properties of homomeric LRRC8A channels in cells: a voltage-dependent conductance activated by low cytoplasmic ionic strength (Kasuya et al., 2018; Syeda et al., 2016).

**Figure 1.**
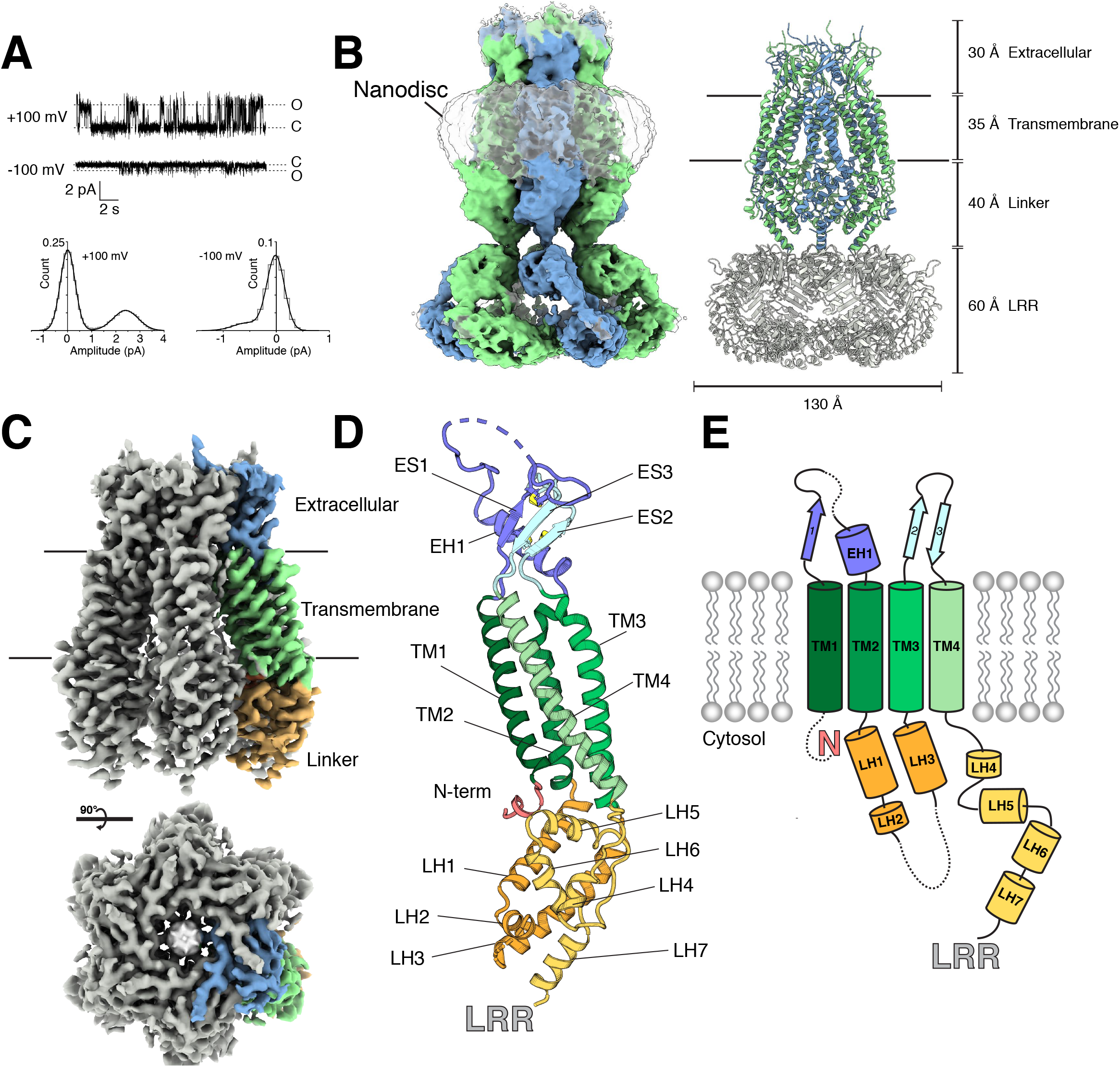
Structure of an LRRC8A-DCPIB complex in lipid nanodiscs.

(A) Representative single channel recording from an excised patch containing purified LRRC8A reconstituted into phosphatidyl choline lipids (P_o_ = 0.3,Y=24 pS at +100mV; P_o_ = 0.24, Y=5.4 pS at −100mV). (B, left) Cryo-EM density of LRRC8A from an unmasked refinement of the constricted state at 4.5 Å resolution and (right) corresponding atomic model viewed from the membrane plane. Individual subunits are alternatingly colored blue and green, nanodisc density is rendered transparent, and LRRs docked into the density are colored gray in the model. Dimensions of the extracellular, transmembrane, linker, and LRR regions are indicated. (C) Cryo-EM density of LRRC8A from a LRR masked refinement of the constricted class at 3.4 Å. A view from the membrane (top) and extracellular space (bottom) are shown. One subunit is colored according to region and shown in (C) within the density, (D) as an isolated model with helices and N-terminus labeled and extracellular domain disulfide bonds depicted in yellow, and (E) as a cartoon.

Initial electron micrographs of LRRC8A solubilized in detergent displayed heterogeneity between particles, especially in the cytoplasmic LRRs (Figure 1-supplement 2). Inspired by recent reports of differences between structures of membrane proteins determined in lipidic and detergent environments (Dang et al., 2017; Jin et al., 2017; Schuler et al., 2013), we pursued high resolution structures of LRRC8A reconstituted in lipid nanodiscs. To this end, LRRC8A was solubilized and purified in detergent, exchanged into lipid nanodiscs formed by the membrane scaffold protein MSP1E3D1 and phosphatidylcholine lipids (POPC, 1-palmitoyl-2-oleoyl-*sn*-glycero-3-phosphocholine), and vitrified in the presence of the LRRC8 inhibitor DCPIB (Figure 1-supplement 1) (Ritchie et al., 2009).

Figure 1B depicts an unmasked refinement of LRRC8A-DCPIB in nanodiscs and accompanying model. LRRC8A forms a 565 kDa, homohexameric channel with each monomer consisting of (from extracellular to intracellular side) an extracellular cap, four transmembrane spanning helices TM1-4, a linker region, and a LRR domain (Figure 1B-E). Initial reconstructions showed high-resolution features in the extracellular cap and transmembrane spanning regions, but less well resolved and blurred features in the linkers and LRR regions (Figure 1-supplement 3). We therefore asked whether distinct conformational states of LRRC8A could be distinguished. Focused classifications on the linker region indeed resolved two channel conformations: a constricted class and an expanded class (Figure 1-supplements 3-5). Significant structural heterogeneity remains in the LRR region within each class. Masking out the LRRs and applying six-fold (C6) symmetry resulted in the highest resolution reconstructions: 3.4 Å resolution for the constricted (Figure 1C) and 3.6 Å resolution for the expanded class, respectively. These reconstructions enabled *de novo* model building and refinement for all regions except for the LRR domain (Figure 1 and Figure 1-supplements 6 and 7), which was instead rigid-body docked into the density from unmasked reconstructions using the high-resolution crystal structure (PDB ID: 6FNW) of this region as a model (Deneka et al., 2018).

### DCPIB inhibits LRRC8A through a cork-in-bottle mechanism

The extracellular cap of LRRC8A is formed by the TM1-TM2 and TM3-TM4 connections from each chain, which pack into a compact domain consisting of a three stranded beta sheet (ES1-3) and short alpha helix EH1 stabilized by three intra-chain disulfide bonds (Figure 1D,E). The subunits associate to form a tube extending ~25 Å above the membrane surface with the inner surface of the tube formed by helix EH1. The N-terminal arginine residue of this helix (R103) projects in towards the conduction axis to form the highly electropositive selectivity filter (Deneka et al., 2018) of the anion-selective channel (Figure 2). The electropositive character in this region is also contributed by the helical dipole of EH1 projecting towards the conduction pathway.

**Figure 2.**
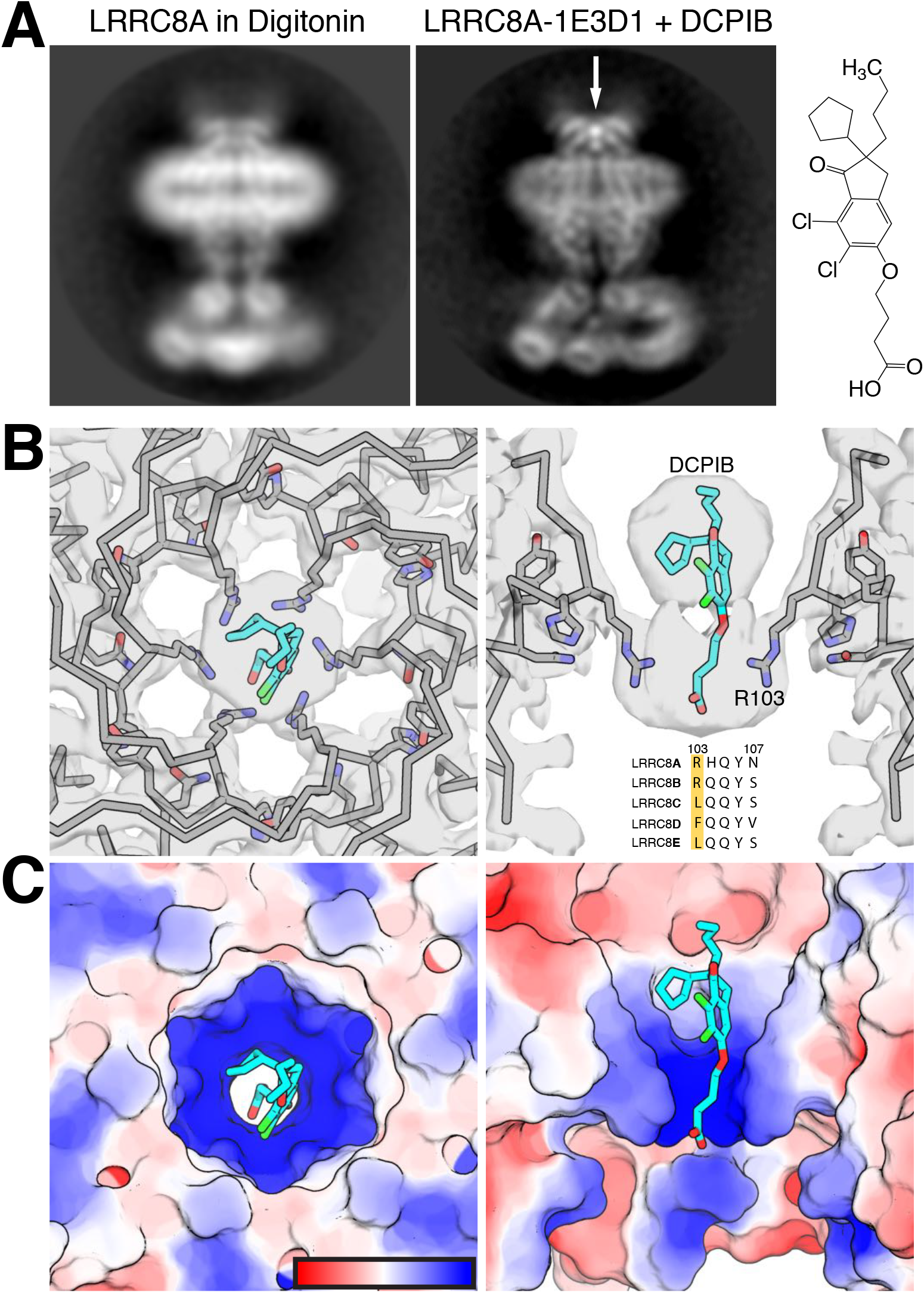
DCPIB inhibitor binding site. (A) Representative side-view two-dimensional class averages of LRRC8A (left) solubilized in digitonin and (middle) reconstituted in MSP1E3D1 lipid nanodiscs and complexed with DCPIB. An arrow highlights the bi-lobed feature present in LRRC8A-DCPIB in nanodiscs, but absent in LRRC8A-digitonin, corresponding to DCPIB in the channel selectivity filter. (right) The chemical structure of DCPIB. (B) View of the selectivity filter with bound DCPIB from (left) the extracellular solution top view and (right) the membrane plane side view. The atomic model is shown as ribbons and sticks within the cryo-EM density with the two front and two rear subunits removed in the side view for clarity. Nitrogens are colored blue, oxygens red, chlorines green, protein carbons gray, and DCPIB carbons teal. Alignment of the residues surrounding the selectivity filter for LRRC8 paralogs is shown below the drug density. (C) Views of the DCPIB binding site as in (B), but with the atomic surface colored by electrostatic potential from electronegative red (−5 k_b_Te_c_^-1^) to electropositive blue (+5 k Te_c_^-1^), with the color scale drawn on the left panel.

The most prominent density in the reconstruction, visible even in two-dimensional class averages, is a champagne cork-shaped feature found within and immediately above the R103 ring (Figure 2A). We attribute this bi-lobed density, not observed in apo-LRRC8A reconstructions, to the DCPIB inhibitor based on size, shape, and chemical considerations (Figure 2A). Attempts to visualize a single binding pose for DCPIB by focused refinement, symmetry-free refinement, or symmetry expansion were unsuccessful. Therefore, the density in our maps represents a six-fold average of positions adopted by DCPIB in different particles (Figure 2B,C). The density is well fit with the predominantly hydrophobic indane, butyl, and cyclopentyl constituents in the larger lobe extracellular to the R103 ring and the negatively charged butanoic acid group in the smaller lobe adjacent to and below the guanidinium groups of the R103 ring. In this way, the electronegative portion of DCPIB interacts favorably with the electropositive arginine side chains (Figure 2C). The connection between the two lobes corresponds to the ester linkage between the indane and butanoic acid. The weaker density for this region is presumably due to it adopting different conformations in different particles and the imposed six-fold symmetry.

The structure of DCPIB bound to LRRC8A provides a simple explanation for the mechanism of drug block: DCPIB acts like a cork to plug the mouth of the channel. The R103 ring electrostatically interacts with the DCPIB carboxylic acid while the bulky hydrophobic end of the drug is sterically too large to pass through the selectivity filter. Ion conduction is thus prevented by an obstructed selectivity filter. DCPIB universally blocks LRRC8 currents even though the key amino acids for the channel-drug interaction (R103 and surrounding residues) are not conserved in LRRC8A paralogs B-E (Figure 2B, right). However, as all functional LRRC8 channels contain LRRC8A subunits, heteromer block by DCPIB is likely due to similar drug-LRRC8A interactions that position the hydrophobic moiety above a selectivity filter too narrow for drug passage.

### Constricted and expanded structures suggest a mechanism to relay conformational changes between an ionic strength sensor and channel gate

Focused classification on the linker region and lower portion of the TMs resolved a constricted and an expanded state of LRRC8A (Figure 3A and Figure 1-supplement 4). The structure of the extracellular cap, DCPIB-bound selectivity filter, and extracellular halves of the TM region are essentially indistinguishable between the two structures (mean Cα r.m.s.d. <0.5 Å). Alignment of the two structures by the extracellular cap shows that the differences are confined to the cytoplasmic halves of the TMs, linker regions, and LRRs. The differences can be approximated as a rigid body displacement of the lower portion of the channel about hinges approximately half way through each transmembrane helix. This generates displacements of ~2-4 Å in the cytoplasmic ends of the transmembrane helices, the linkers, and at the cytoplasmic connection between the linkers and LRRs (Figure 3B-D). As a consequence of these conformational changes, the channel cavity is markedly larger in the expanded state: it dilates approximately 4 Å including at the N-terminal helix at Pro15 (Figure 3B and Movies 1 and 2).

**Figure 3.**
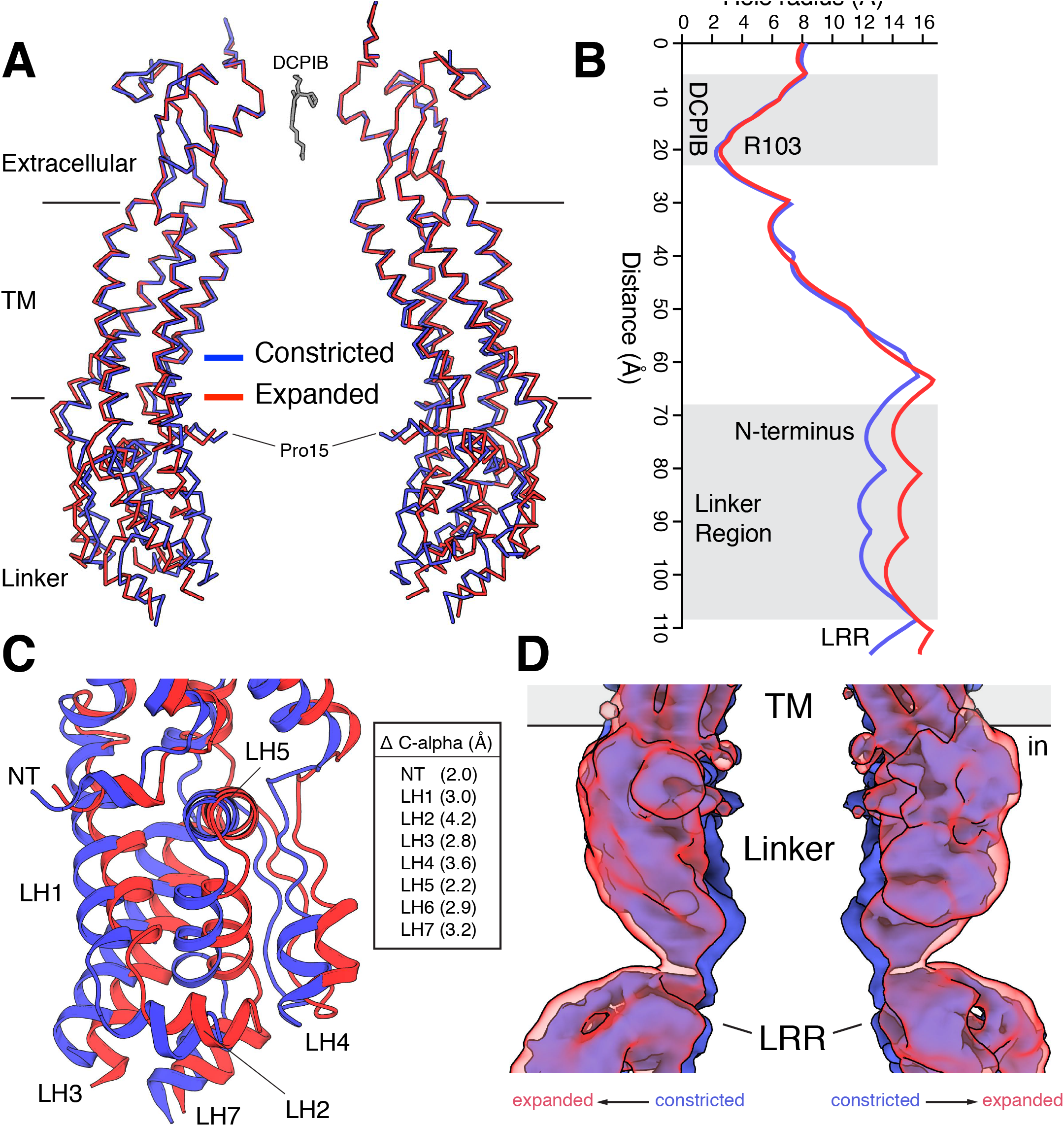
Constricted and expanded LRRC8A structures. (A) Overlay of constricted (blue) and expanded (red) structures of LRRC8A viewed from the membrane with two opposing subunits shown for each structure in ribbon representation. DCPIB is shown in grey sticks. Proline 15, the final modeled residue at the N-terminus, is labeled. (B) Comparison of the pore radius along the conduction axis colored as in (A). (C) Close-up view of the structure overlay at the linker region with models drawn as cartoons. Helices are labeled and distances between C**α** positions at the following positions are indicated: NT, R18; LH1, K162; LH2, T170; LH3, K249; LH4, I356; LH5, S379; LH6, S387; LH7, E399. (D) Overlay of unmasked cryo-EM maps from constricted and expanded particles showing correlated movement of the linker region and membrane-proximal LRR region. Density within 5Å of two opposing chains is shown.

Might these differences be related to gating conformational changes? Gating in LRRC8 and related connexin and innexin channels is thought to involve the N-terminus prior to the beginning of TM1 (Kefauver et al., 2018a; Oshima, 2014; Zhou et al., 2018), whereas sensing of internal ionic strength is thought to occur in the cytoplasmic LRRs or connection between linker helices 2 and 3 (Syeda et al., 2016). Notably, the conformational changes we observe between constricted and expanded states couple these presumed sensing and gating elements. The extreme N-terminus of LRRC8A (amino acids 1-14) is not resolved in our structures (or in detergent-solubilized LRRC8A structures (Deneka et al., 2018; Kasuya et al., 2018; Kefauver et al., 2018a)). However, amino acids 15-21 are visible and form a helical extension of TM1 that projects towards the center of the channel (Figure 3A and 3B). This region is coupled through the linkers to the LRRs via an elbow formed by the N-terminal helix, TM1, and LH5 (Figure 3C). A network of hydrophobic contacts including L20 and L376 rigidifies the connection between the LRRs and N-terminus. Movement of cytoplasmic regions, perhaps as a consequence reduced ionic strength, can thus be coupled to the N-terminal gating region and an expansion of the channel pore in a concerted fashion.

### Channel symmetry and heterogeneity of LRRs in nanodiscs

Recent structures of LRRC8A in digitonin showed an unexpected three-fold symmetric trimer of dimers arrangement with pairs of LRRC8A subunits forming an asymmetric unit of the channel (Deneka et al., 2018; Kasuya et al., 2018; Kefauver et al., 2018a). Strikingly, LRRC8A in lipid nanodiscs does not display this arrangement. Instead, we observe six-fold symmetric channels with conformationally heterogeneous LRRs.

Structural heterogeneity in LRRs of particles in lipid nanodiscs is readily apparent in 2D class averages (Figure 4A, Movies 3 and 4). Side views of the particles constituting both constricted and expanded classes display a range of LRR positions: LRRs are closely packed underneath the linker region in some classes and splayed laterally away from the conduction axis in others. As expected from the two-dimensional classes, three-dimensional reconstructions with or without symmetry enforced, or using masking strategies to isolate the LRRs, generated low-resolution features for this region. We conclude that LRRs of LRRC8A-DCPIB in lipid nanodiscs can access a large conformational space. Still, the average position of membrane proximal region of the LRRs adopts a more constricted or expanded position in the corresponding structure class, about which there appears to be similar heterogeneity (Figures 3D and Figure 4A).

**Figure 4.**
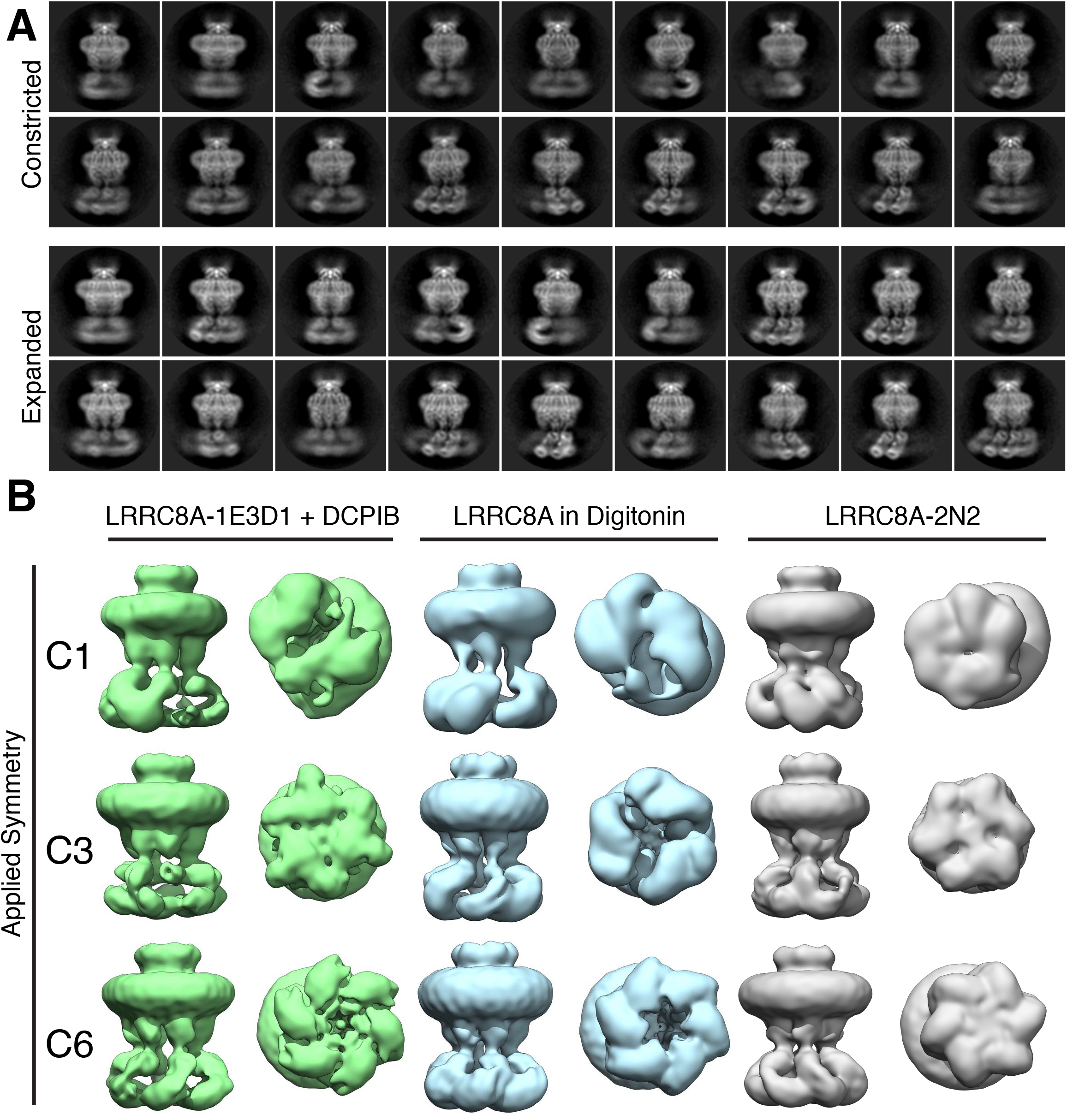
LRR position and channel symmetry differences in lipid and detergent environments. (A) Side-views of two-dimensional class averages from the (top) constricted and (bottom) expanded particle classes illustrating variation in LRR position. Also see Movies 3 and 4. (B) Symmetry comparison of LRRC8A in (left) MSP1E3D1+DCPIB, (middle) digitonin, or (right) MSP2N2 (displayed at a 0.015 threshold). Selected classes from three-dimensional classing jobs with the indicated symmetry are shown from the side (membrane) or bottom (cytoplasm).

We used three approaches to assess the symmetry of LRRC8A in lipid nanodiscs. First, we subjected LRRC8A-nanodisc particle stacks (generated without imposing symmetry) to classification without symmetry (C1), with three-fold (C3), or with six-fold (C6) symmetry imposed. The results are compared in Figure 4B and Figure 4-supplements 1 and 2. Reconstructions without imposed symmetry have six-fold symmetric features including in the linker region where individual subunits are oriented and spaced uniformly. As expected, reconstructions with three- and six-fold symmetry imposed recapitulate these features with improved definition and resolution. As a second approach, we subjected the final LRRC8A-nanodisc particle stacks used for high-resolution reconstructions (generated with C6-symmetry imposed and LRRs masked) to refinements with C1, C3, or C6 symmetry imposed. C1 reconstructions display features with six-fold symmetry. Imposed C3 and C6 symmetry recapitulate these map features with improved detail. As a third approach, we asked whether references made from reported C3-symmetric structures of LRRC8A in digitonin could be used to identify a class of C3 symmetric particles in nanodiscs. This approach failed to generate high-resolution reconstructions with or without masking of the LRRs. We therefore conclude that LRRC8A-DCPIB in nanodiscs is six-fold symmetric outside of the heterogeneous LRRs.

To test whether differences between nanodisc and digitonin structures are related to different hydrophobic environments or other factors, we imaged apo-LRRC8A in digitonin and in MSP2N2 nanodiscs and performed analogous analyses (Figure 4B and Figure 4-supplements 1 and 2). Consistent with published structures (Deneka et al., 2018; Kasuya et al., 2018; Kefauver et al., 2018a), approximately one-third of the digitonin-solubilized apo-LRRC8A particles contribute to a class with compact LRRs. Reconstructions without imposed symmetry have three-fold symmetric features (i.e. a trimer-of-dimers arrangement) that are clarified with C3 symmetry and obscured with C6 symmetry imposed. In contrast, reconstructions of apo-LRRC8A in MSP2N2 nanodiscs are similar to LRRC8A-DCPIB in MSP1E3D1 nanodiscs, albeit at lower resolution. Thus, the presence of DCPIB, choice of scaffold protein, or expression host are unlikely to generate structural and symmetry differences in LRRC8A structures. Instead, we conclude that the hydrophobic environment surrounding LRRC8A is a key determinant of channel structure: digitonin micelles promote three-fold channel symmetry and compact LRRs while lipid bilayers promote six-fold symmetric channels with asymmetric and conformationally heterogeneous LRRs.

How might the differences between digitonin and lipid bilayers influence the structure of LRRC8A channels? A conspicuous feature of LRRC8A is the presence of gaps between subunits that create lipid-facing crevasses in the channel surface. In lipid nanodiscs, a large lower gap and a smaller upper gap are separated by a constriction made by amino acids Leu131 and Phe324. The upper gap is filled with three well-defined tubular features that are modeled as partial POPC lipid hydrocarbon acyl chains (Figure 5). The presence of these ordered acyl chains seals the upper gap in the channel surface and may act as a “glue” that connects adjacent subunits and stabilizes the upper transmembrane domain against movements in channel linkers and LRRs. In the C3-symmetric LRRC8A structures in digitonin (Kasuya et al., 2018; Kefauver et al., 2018a), there are two distinct subunit interfaces – one with a narrow separation and one with a wider separation between neighboring chains (Figure 6 and Movies 5 and 6). This results in a striking rearrangement of the lipid binding site. In nanodiscs, the upper gap tapers to a tunnel ideally sized to surround the central bound acyl chain (5-6 Å in diameter). In detergent, in contrast, this gap becomes either too large to optimally accommodate an acyl chain (6.9-7.3Å diameter in the wide interface) or too small for one to fit (4.1 Å diameter in the narrow interface). Consistently, density for lipids is not observed in LRRC8A-digitonin reconstructions. In one map, a feature consistent with a digitonin molecule is wedged between subunits near the wide-interface upper gap instead (Figure 6C) (Kefauver et al., 2018b). This suggests an explanation for the differences between LRRC8A structures: when lipid surrounding the channel is removed, the channel adapts from six evenly spaced lipid-filled gaps to three smaller gaps incompatible with lipid binding and three larger gaps perhaps filled with digitonin. These changes at subunit interfaces in the transmembrane region are then propagated through the structure to influence overall channel symmetry and LRR position.

**Figure 5.**
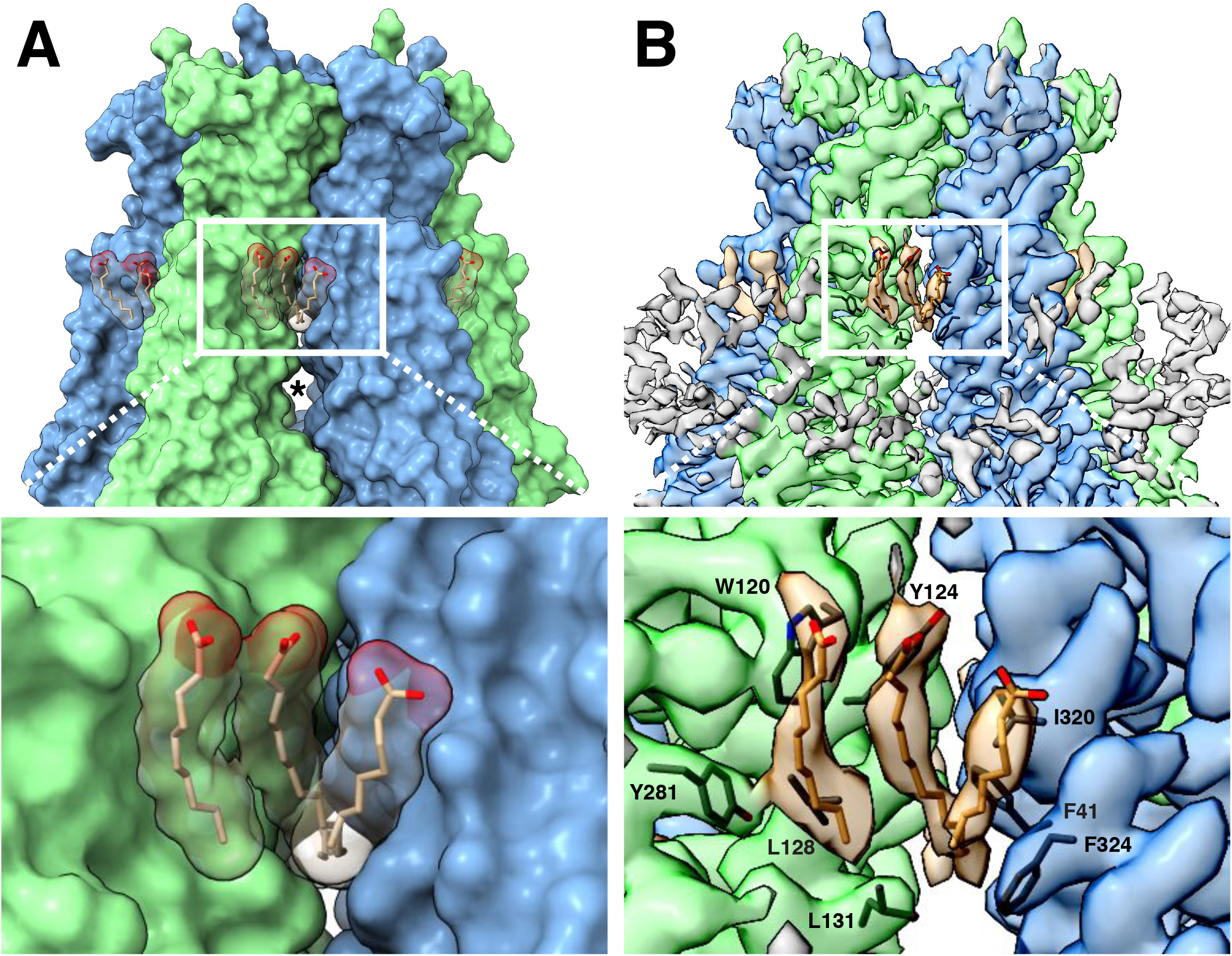
LRRC8A-lipid interactions. (A, above) Surface representation of the constricted LRRC8A class viewed from the membrane with docked POPC lipid chains depicted in stick (tan) and space-filling (transparent) representations. The upper gap between subunits is filled by lipid density and an asterisk marks the larger lower gap. (Below) a zoomed-in view on the upper gap and bound lipid. (B, above) The corresponding cryo-EM density viewed as in (A), with lipid and the nearby hydrophobic amino-acid side chains shown in stick form. (Below), a zoomed-in view with the amino acids of the hydrophobic pocket labeled.

**Figure 6.**
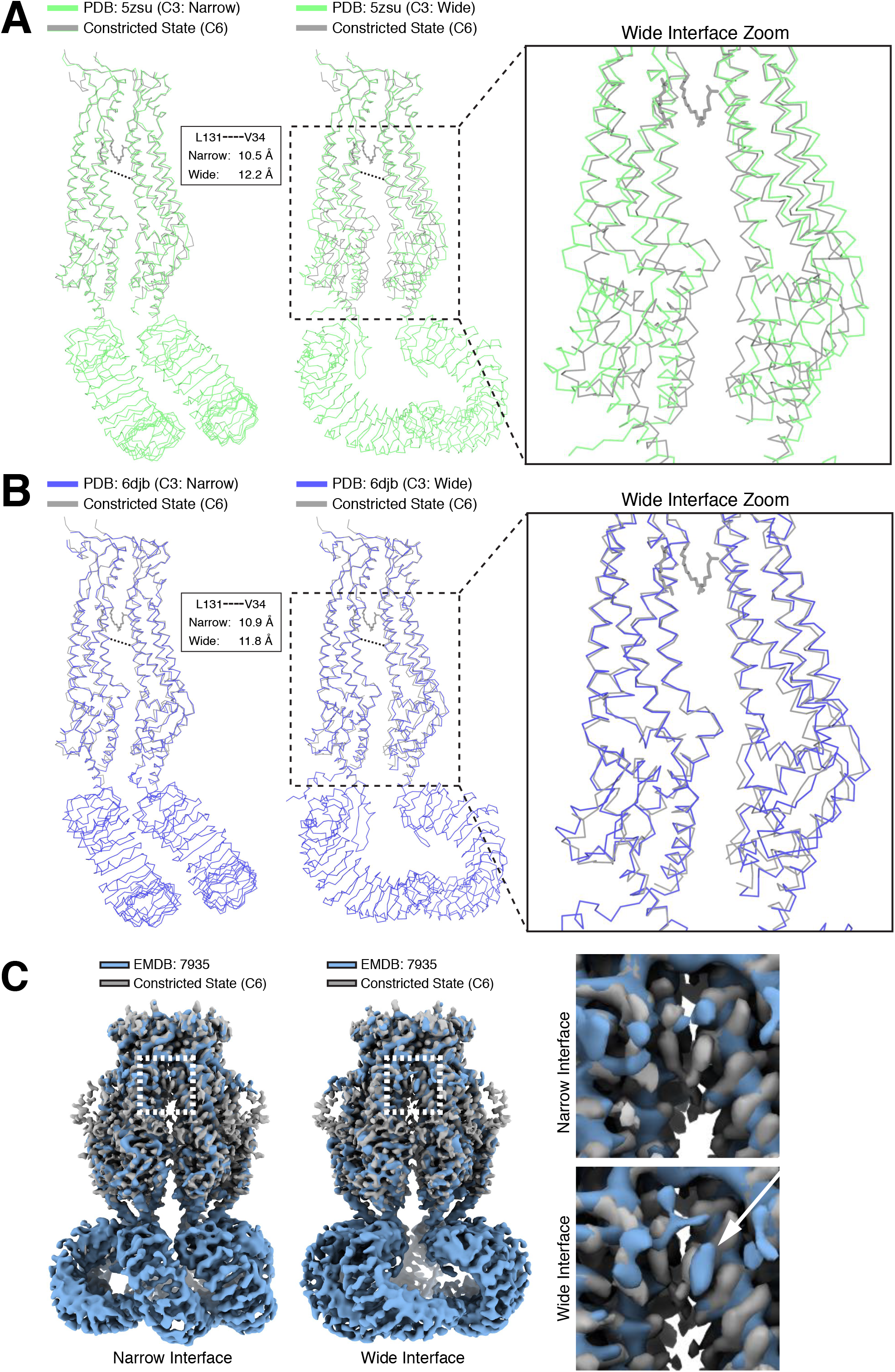
Differences in LRRC8A structures solved in lipid bilayers and detergent micelles. Overlays of extracellular domain-aligned models of the constricted state structure determined in lipid nanodiscs (gray) and the (left) narrow and (middle) wide subunit interfaces from structures determined in digitonin: (A, PDB: 5zsu, green (Kasuya et al., 2018)), (B, PDB: 6djb, blue (Kefauver et al., 2018a)). Boxed and zoomed in regions (right) illustrate the expansion in subunits of the wide interfaces extending to the LRR region. Distances between V34 and L131 C**α**s in detergent-solubilized structrues are indicated. (C) Aligned density between the six-fold symmetric constricted state in nanodiscs (gray) and LRRC8A in digitonin (Blue, EMDB: 7935, (Kefauver et al., 2018a)). (left) View from the membrane of the narrow C3 subunit interface. (middle) View from the membrane of the wide C3 subunit interface. Boxed region and zoomed-in panels (right) show the upper gap filled with lipid density in nanodiscs or an extra density consistent with digitonin in the wide interface marked by a white arrow.

### Discussion

The nanodisc-embedded LRRC8A-DCPIB structures presented here provide the first insights into drug block of LRRC8 channels or related connexin and innexin channels. DCPIB binding in the channel selectivity filter is largely dictated by size and electrostatic complementarity of the small molecule and channel. The relatively modest (low micromolar) affinity of DCPIB for LRRC8 channels (Decher et al., 2001) is probably due to the small protein-DCPIB interface and lack of significant structural rearrangement in the selectivity filter upon binding. These characteristics are also consistent with the broad target profile of DCPIB; in addition to inhibiting LRRC8 homomeric and heteromeric channels, it also inhibits connexin hemichannels (Bowens et al., 2013). The structures reported here provide a platform for future structure-based design of inhibitors with higher affinity or more restricted target range. Improved pharmacology would likely help to both refine reported roles and define new physiological roles for LRRC8 channels.

The correlated changes in the N-terminal region, linker, and LRRs in the constricted and expanded states suggest a mechanism to transmit information from the ionic strength sensor to the channel gate. We speculate that the expanded state represents an intermediate on the path to a fully conductive channel. The 4 Å increase in channel cavity diameter at the position of the unresolved N-terminus represents a significant expansion, comparable in size to the selectivity filter opening and approaching the diameter of a solvated chloride ion. Still, additional conformational changes are likely required to fully open the channel. The particles are vitrified in an ionic strength that would correspond to closed or inactivated channels (150 mM KCl) and the N-terminal fourteen amino acids that project into the channel cavity and are implicated in channel gating are unresolved due to lack of a consistent structure between particles. One intriguing possibility is that a dynamic N-terminal region creates an entropic barrier to ion passage in nonconductive channels and gating by low ionic strength involves conformational changes that order the N-terminus to create a favorable environment for anion conduction.

Interestingly, both the symmetry changes and conformational flexibility of the LRRs are reminiscent of the unrelated bacterial CorA channel (Deneka et al., 2018; Kefauver et al., 2018a; Matthies et al., 2016). In CorA, loss of intracellular Mg^2+^ binding allows intracellular domains to move outward and adopt an asymmetric conformation that promotes conduction. In LRRC8A, low ionic strength may analogously result in loss of interactions between cytoplasmic LRRs and/or linker helix 2-3 connections that otherwise promote a compact structure and constricted pore. This could allow expansion in cytoplasmic regions that is relayed through the linkers to dilate the channel and influence the conformation of the N-terminal gating region. Future studies that define conductive LRRC8 structures and correlate channel motions with functional states will help to elucidate the ionic-strength sensing mechanism of this dynamic and prolific vertebrate channel family.

**Table 1.** Cryo-EM data collection, processing, refinement, and modeling data for LRRC8A-DCPIB in MSP1E3D1 nanodiscs.

| **Data collection** | **Sloan-Kettering** | **NYSBC** |
| --- | --- | --- |
| Movie # | 2482 | 2110 |
| Magnification | 22,500x | 22,500x |
| Voltage (kV) | 300 | 300 |
| Electron exposure (e<sup>-</sup>/Å<sup>2</sup>) | 60.8 | 55.6 |
| Defocus range (μm) | -1.2~-2.5 | -1.2~-2.5 |
| Super resolution pixel size (Å) | 0.544 | 0.536 |
| Fourier cropped pixel size (Å) | 1.088 | 1.088 |
| **Processing** | **Class 1 (constricted)** | **Class 2 (expanded)** |
| Symmetry imposed | C6 | C6 |
| Initial particle images (no.) | 752,736 | 752,736 |
| Final particle images (no.) | 25,153 | 35,435 |
| Map resolution (Å) / FSC threshold | 3.39 / 0.143 | 3.63 / 0.143 |
| **Refinement** |  |  |
| Model resolution (Å) | 3.52 / 3.32 | 3.81 / 3.47 |
| FSC threshold | 0.50 / 0.143 | 0.50 / 0.143 |
| Map-sharpening *B*factor (Å<sup>2</sup>) | -44.538 | -134.6 |
| Ligands | 19 | 19 |
| Mean *B* factors (Å<sup>2</sup>) |  |  |
| Protein | 87.73 | 52.17 |
| Ligand | 65.05 | 27.6 |
| R.m.s. deviations |  |  |
| Bond lengths (Å) | 0.007 | 0.005 |
| Bond angles (°) | 0.775 | 0.823 |
| Validation |  |  |
| MolProbity score | 1.74 | 1.94 |
| Clashscore | 3.34 | 4.12 |
| Poor rotamers (%) | 3.09 | 3.17 |
| EMRinger score | 2.72 | 2.2 |
| Ramachandran plot |  |  |
| Favored (%) | 96.42 | 94.67 |
| Allowed (%) | 3.58 | 5.33 |
| Disallowed (%) | 0 | 0 |

**Table 2.** Cryo-EM data collection information for the digitonin and MSP2N2 datasets used in Figure 2A and Figure 4.

| Data collection | Digitonin 70 mM KCl | Digitonin 150 mM KCl | Digitonin 600 mM KCl | MSP2N2 |
| --- | --- | --- | --- | --- |
| Movie # | 2550 | 1445 | 1464 | 1786 |
| Magnification | 22500x | 22,500x | 22,500x | 22,500x |
| Voltage (kV) | 300 | 300 | 300 | 300 |
| Electron exposure (e<sup>-</sup>/Å<sup>2</sup>) | 60.8 | 55.6 | 55.6 | 70.3 |
| Defocus range (μm) | -1.2~-2.5 | -1.2~-2.5 | -1.2~-2.5 | -1.2~-2.5 |
| Super resolution pixel size (Å) | 0.536 | 0.544 | 0.544 | 0.536 |
| Fourier cropped pixel size (Å) | 1.088 | 1.088 | 1.088 | 1.115 |

## Acknowledgements

We thank the members of the Brohawn laboratory for support, input, and critical reading of the manuscript. We gratefully acknowledge Team Brohawn (B. Phillips, H. Falahati, J. Quiroz) and R. Sepela for help with recordings at the 2018 Electrophysiology course at the Marine Biological Institute. We thank E. Montabana for training in grid preparation and sample screening and D. Toso and P. Tobias at the Berkeley Bay Area Cryo-EM facility for help with collection of preliminary data. We thank M. de la Cruz at the Memorial Sloan Kettering Cancer Center Cryo-EM Facility and the staff at the Simons Electron Microscopy Center for help with data collection. SGB is a New York Stem Cell Foundation-Robertson Neuroscience Investigator. This work is supported by the New York Stem Cell Foundation (SGB), NIGMS grant DP2GM123496-01 (SGB), a McKnight Foundation Scholar Award (SGB), a Klingenstein-Simons Foundation Fellowship Award (SGB), NIH-NCI Cancer Center Support Grant (PO CA008748), a Josie Robertson Investigators award (RKH), and the Searle Scholars Program (RKH). Some of this work was performed at the Simons Electron Microscopy Center and National Resource for Automated Molecular Microscopy located at the New York Structural Biology Center, supported by grants from the Simons Foundation (SF349247), NYSTAR, and the NIH National Institute of General Medical Sciences (GM103310) with additional support from Agouron Institute (F00316) and NIH (OD019994).

## Data availability

Final maps of LRRC8A-DCPIB in nanodiscs have been deposited to the Electron Microscopy Data Bank under accession codes XXX (unmasked constricted state), XXX (masked constricted state), XXX (unmasked expanded state), and XXX (masked expanded state). Atomic coordinates have been deposited in the PDB under IDs XXX (constricted state) and XXX (expanded state). Micrographs are deposited with accession codes XXX. All data are available upon request to the corresponding author(s).

## Methods

### Protein Expression

The coding sequence for LRRC8A from *Mus musculus* was codon optimized for *Spodoptera frugiperda* and synthesized (Gen9, Cambridge, MA). The sequence was then cloned into a custom vector based on the pACEBAC1 backbone (MultiBac; Geneva Biotech, Geneva, Switzerland) with an added C-terminal PreScission protease (PPX) cleavage site, linker sequence, superfolder GFP (sfGFP) and 7xHis tag, generating a construct for expression of mmLRRC8A-SNS-LEVLFQGP-SRGGSGAAAGSGSGS-sfGFP-GSS-7xHis. MultiBac cells were used to generate a Bacmid according to manufacturer’s instructions. *Spodoptera frugiperda* (Sf9) cells were cultured in ESF 921 medium (Expression Systems, Davis, CA) and P1 virus was generated from cells transfected with Cellfectin II reagent (Life Technologies, Carlsbad, CA) according to manufacturer’s instructions. P2 virus was then generated by infecting cells at 2 million cells/mL with P1 virus at an MOI ~0.1, with infection monitored by fluorescence of sfGFP-tagged protein and harvested at 72 hours. P3 virus was generated in a similar manner to expand the viral stock. The P3 viral stock was then used to infect 1 L of Sf9 cells at 4 million cells/mL at an MOI ~2–5. At 72 hours, infected cells containing expressed LRRC8A-sfGFP protein were harvested by centrifugation at 2500 x g and frozen at −80°C.

### Protein Purification

Cells from 1 L of culture (~15-20 mL of cell pellet) were thawed in 100 mL of Lysis Buffer containing (in mM) 50 HEPES, 150 KCl, 1 EDTA pH 7.4. Protease inhibitors (Final Concentrations: E64 (1 *μ*M), Pepstatin A (1 *μ*g/mL), Soy Trypsin Inhibitor (10 *μ*g/mL), Benzimidine (1 mM), Aprotinin (1 *μ*g/mL), Leupeptin (1*μ*g/mL), and PMSF (1mM)) were added to the lysis buffer immediately before use. Benzonase (4 *μ*l) was added after cell thaw. Cells were then lysed by sonication and centrifuged at 150,000 x g for 45 minutes. The supernatant was discarded and residual nucleic acid was removed from the top of the membrane pellet using DPBS. Membrane pellets were scooped into a dounce homogenizer containing Extraction Buffer (50 mM HEPES, 150 mM KCl, 1 mM EDTA, 1% n-Dodecyl-β-D-Maltopyranoside (DDM, Anatrace, Maumee, OH), 0.2% Cholesterol Hemisuccinate Tris Salt (CHS, Anatrace) final pH 7.4). A 10%/2% solution of DDM/CHS was dissolved and clarified by bath sonication in 200 mM HEPES pH 8 prior to addition to buffer to the indicated final concentration. Membrane pellets were then homogenized in lysis buffer and this mixture was gently stirred at 4°C for 3 hours. The extraction mixture was centrifuged at 33,000 x g for 45 minutes and the supernatant, containing solubilized membrane protein, was bound to 4 mL of sepharose resin coupled to GFP nanobody for 1 hour at 4°C. The resin was then collected in a column and washed with 10 mL of Buffer 1 (20 mM HEPES, 150 mM KCl, 1 mM EDTA, 0.025% DDM, pH 7.4), 40 mL of Buffer 2 (20 mM HEPES, 500 mM KCl, 1 mM EDTA, 0.025% DDM, pH 7.4), and 10 mL of Buffer 1. The resin was then resuspended in 6 mL of Buffer 1 with 0.5 mg of PPX and rocked gently in the capped column for 2 hours. Cleaved LRRC8A protein was then eluted with an additional 8 mL of Wash Buffer, spin concentrated to ~500 *μ*l with Amicon Ultra spin concentrator 100 kDa cutoff (Millipore), and then loaded onto a Superose 6 Increase column (GE Healthcare, Chicago, IL) on an NGC system (Bio-Rad, Hercules, CA) equilibrated in buffer 1. Peak fractions containing LRRC8A channel were then collected and spin concentrated.

Purification in digitonin was performed analogously with the following modifications. Tris buffer was used instead of HEPES buffer during purification. Washing and cleavage steps were carried out in 0.1% DDM and 0.02% CHS and cleavage was performed overnight. Digitonin (EMD Chemicals Inc., San Diego, CA) Buffer containing 20 mM Tris, (70, 150, or 600) mM KCl, 1 mM EDTA, 0.05% Digitonin (final pH 8) was prepared by dissolving digitonin in buffer at room temperature, cooling buffer to 4°C, and 0.2 *μ*m filtering the buffer to remove insoluble material. Protein was exchanged into digitonin buffer by gel filtration in a Superose 6 Increase column. Fractions containing LRRC8A channel were pooled and spin concentrated.

### Electrophysiology

For proteoliposome patching experiments, we incorporated protein into lipid and generated proteoliposome blisters for patch recordings using dehydration and rehydration as described previously (Brohawn et al., 2014; Del Mármol et al., 2018) with the following modifications. LRRC8A was first purified into Column Buffer with DDM/CHS at 0.025%/0.005%. Protein was then exchanged into lipid with the addition of Biobeads SM2 and an overnight incubation. Dried proteoliposomes in a dish were rehydrated overnight in a buffer containing 10 mM HEPES, 70 mM KCl, pH 7.4. The next day, the dish was filled with a bath solution containing 10 mM HEPES, 20 mM MgCl_2_, 30 mM KCl pH 7.4. The pipette solution was 10 mM HEPES, 70 mM KCl, pH 7.4. All experiments were conducted at room temperature.

### Nanodisc Formation

Freshly purified LRRC8A from gel filtration in Buffer 1 was reconstituted into MSP1E3D1 nanodiscs with POPC lipid (Avanti, Alabaster, Alabama) at a final molar ratio of 1:2.5:250 (Monomer Ratio: LRRC8A, MSP1E3D1, POPC). First, solubilized lipid in Column Buffer (20 mM HEPES, 150 mM KCl, 1 mM EDTA pH 7.4) was mixed with additional DDM detergent, Column Buffer, and LRRC8A. This mix was mixed at 4°C for 30 minutes before addition of purified MSP1E3D1. This addition brought the final concentrations to approximately 10 *μ*M LRRC8A, 25 *μ*M MSP1E3D1, 2.5 mM POPC, and 4 mM DDM in Column Buffer. The solution with MSP1E3D1 was mixed at 4°C for 30 minutes before addition of 160 mg of Biobeads SM2 (Bio-Rad). Biobeads (washed into methanol, water, and then Column Buffer) were weighed with liquid removed by P1000 tip (Damp weight). This mix was incubated at 4°C for 30 minutes before addition of another 160 mg of Biobeads (final 320 mg of Biobeads per mL). This final mixture was then mixed at 4°C overnight (~ 12 hours). Supernatant was cleared of beads by letting large beads settle and by 0.2 *μ*m filtering. Sample was spun for 5 minutes at 21,000 x g before loading onto a Superose 6 column in Column Buffer. Peak fractions corresponding to LRRC8A in MSP1E3D1 were collected, 100 kDa cutoff spin concentrated, and then re-run on the Superose 6. The fractions corresponding to the center of the peak were then pooled and concentrated prior to grid preparation. For MSP2N2 nanodiscs, reconstitution was carried out with a molar ratio of 1:2:300 (Monomer Ratio: LRRC8A, MSP2N2, Soy Polar Extract (Avanti)). The final 1 mL reaction mix contained 5 *μ*M LRRC8A monomer, 10 *μ*M MSP2N2, 1.5 mM Soy Polar, ~2.6 mM DDM, and 200 mg of Biobeads in Column Buffer. Column purification was performed similarly to the MSP1E3D1 preparation.

### Grid Preparation

For the MSP1E3D1 nanodisc samples, 100 *μ*M of DCPIB (Tocris, Bristol, UK) was added to sample to give a final concentration of 0.8 mg/mL LRRC8A-MSP1E3D1. DCPIB was allowed to equilibrate and bind complex on ice for 1 hour prior to freezing grids. Sample with drug was cleared by a 5 minute 21,000 x g spin prior to grid making. For freezing grids, a 3 *μ*l drop of protein was applied to freshly glow discharged Holey Carbon, 400 mesh R 1.2/1.3 gold grids (Quantifoil, Großlöbichau, Germany). A FEI Vitrobot Mark IV (ThermoFisher Scientific) was utilized with 22°C, 100% humidity, 1 blot force, and a 3 second blot time, before plunge freezing in liquid ethane. Grids were then clipped in autoloader cartridges (FEI, Hillsboro, Oregon) and shipped in a dry shipper for data collection. The MSP2N2 sample was frozen without drug at 0.8 mg/mL with the same conditions as the MSP1E3D1 grids.

For the digitonin sample, purified protein was shipped in a refrigerated container. The next day, for freezing, a 3 *μ*l drop of protein was applied to a freshly glow discharged Holey Carbon, 400 mesh R 1.2/1.3 gold grid (Quantifoil). A FEI Vitrobot Mark IV (ThermoFisher Scientific) was utilized with 22°C, 100% humidity, 1 blot force, and a 5 second blot time, before plunge freezing in liquid ethane. Grids were then clipped and used for data collection.

### Cryo-EM data acquisition

For the digitonin-solublized channels in 150 and 600 mM KCl, grids were transferred to an FEI Titan Krios cryo-EM operated at an acceleration voltage of 300 kV. Images were recorded in an automated fashion with SerialEM (Mastronarde, 2005) with a defocus range of −1.2 ~ −2.5 *μ*m over 8 seconds as 40 subframes with a Gatan K2 direct electron detector in super-resolution mode with a super-resolution pixel size of 0.544 Å. The electron dose was 8 e^-^/pixel/s at the detector level and total accumulated dose was 54 e^-^/Å^2^. For the digitonin-solublized channels in 70 mM KCl, grids were transferred to an FEI Titan Krios cryo-EM operated at an acceleration voltage of 300 kV. Images were recorded in an automated fashion with Leginon (Suloway et al., 2005) with a defocus range of −1.2 ~ −2.5 *μ*m over 8 seconds as 40 subframes with a Gatan K2 direct electron detector in super-resolution mode with a super-resolution pixel size of 0.536 Å. The electron dose was 9 e^-^/pixel/s at the detector level and total accumulated dose was 55.6 e^-^/Å^2^. For MSP2N2 nanodisc-reconstituted samples, grids were transferred to an FEI Titan Krios cryo-EM operated at an acceleration voltage of 300 kV. Images were recorded in an automated fashion with Leginon with a defocus range of −1.2 ~ −2.5 *μ*m over 10 seconds as 40 subframes with a Gatan K2 direct electron detector in super-resolution mode with a super-resolution pixel size of 0.536 Å.

The MSP1E3D1 nanodisc-reconstituted samples were recorded in two sessions. For the first session, grids were transferred to an FEI Titan Krios cryo-EM operated at an acceleration voltage of 300 kV. Images were recorded in an automated fashion with SerialEM with a defocus range of −1.2 ~ −2.5 *μ*m over 8 seconds as 40 subframes with a Gatan K2 direct electron detector in super-resolution mode with a super-resolution pixel size of 0.544 Å. The electron dose was 9 e^-^/pixel/s at the detector level and total accumulated dose was 60.8 e^-^/Å^2^. For the second session, grids were transferred to an FEI Titan Krios cryo-EM operated at an acceleration voltage of 300 kV. Images were recorded in an automated fashion with Leginon with a defocus range of −1.2 ~ −2.5 *μ*m over 8 seconds as 40 subframes with a Gatan K2 direct electron detector in super-resolution mode with a super-resolution pixel size of 0.536 Å. The electron dose was 9 e^-^/pixel/s at the detector level and total accumulated dose was 55.6 e^-^/Å^2^.

### Cryo-EM Data Processing

Processing was carried out using Relion 3.0 (Zivanov et al., 2018b), (Zivanov et al., 2018a). Movies were gain and motion corrected with the Relion MotionCor2 package (standalone MotionCor2 for the digitonin and MSP2N2 datasets) (Zheng et al., 2017), and the datasets were each binned to 1.088 Å/pixel. Ctf estimation was performed with Gctf 1.06. For particle picking, 1000-2000 particles were picked manually to generate references for autopicking. For the MSP1E3D1 datasets, 2x particle binning was performed at extraction for initial particle cleanup, before re-extraction to 1.088 Å/pixel for final classification and refinement. For the other datasets, particles were 2x binned for all processing steps except final 2D comparisons.

For the contracted and expanded LRRC8A states, we first noted differences in the linker region during initial full particle classing and blurred helical density in this region during full particle refinement. We therefore used a mask encompassing the linker region and the bottom of the transmembrane helices during classification without angular sampling to separate particles in these two classes. To obtain high-resolution particles for the final reconstruction, masking out the LRR region was crucial. For the final particle sets, we also performed unmasked refinement, which generated the full particle map shown in Figure 1 for the contracted state. The full particles for the expanded state were consistently more difficult to refine to high resolution and also saw no benefit from Ctf Refinement, potentially due to their additional LRR heterogeneity.

For symmetry testing on the digitonin, MSP2N2, and MSP1E3D1 datasets, 2x binned particles (2.176 Å/pixel or 2.298 Å/pixel for MSP2N2) were first cleaned using 2D classification and 3D Classification using C1 symmetry with the same Gaussian filtered initial reference. For final symmetry analysis, three 3D Classifications were performed using the same reference and C1, C3, and C6 symmetry operations. For further comparison of digitonin and MSP2N2 particles, the best C1 classes were then used for final refinements in C3 and C6.

Overall resolution was estimated using Relion 3.0 and Phenix.mtriage. Local resolution was calculated using Relion. For a detailed pipeline see Figure 1-supplements 3-7.

### Modeling and Refinement

Cryo-EM maps were sharpened using Phenix.autosharpen. The structures were modeled ab inito in Coot for all regions outside of the LRRs and refined in real space using Phenix.real_space_refine implementing Ramachandran and NCS restraints. Restraints for DCPIB and POPC ligands were generated using Phenix.elbow from SMILES string inputs and optimized with the eLBOW AM1 QM method. Validation tools in Phenix, EMRinger (Barad et al., 2015), and Molprobity were used to guide two subsequent rounds of iterative manual adjustment in Coot and refinement in Phenix. For cross-validation, atoms in the final model were randomly displaced up to 0.5 Å and refined against one half-map (‘work’). FSC curves were then calculated between the refined model and each half-map (‘work’ and ‘free’) using Phenix.mtriage. The absence of significant differences between the FSC curves is indicates the model was not overfit to the original map. Superpositions with published LRRC8A-detergent structures, which were not used as guides during model building, also demonstrate good overall correspondence aside from symmetry and conformational changes. For illustration of average LRR position in Figure 1, the 1.8 Å crystal structure from PDB 6FNW was docked as a rigid body into unmasked maps using Phenix. Channel cavity measurements were made with HOLE implemented in Coot. Electrostastic potential was calculated using APBS-PDB2PQR (Dolinsky et al., 2004). Figures were prepared using PyMOL, Chimera, ChimeraX, Fiji, Prism, and Adobe Photoshop and Illustrator software.

---

**Figure 1-Supplement 1.**
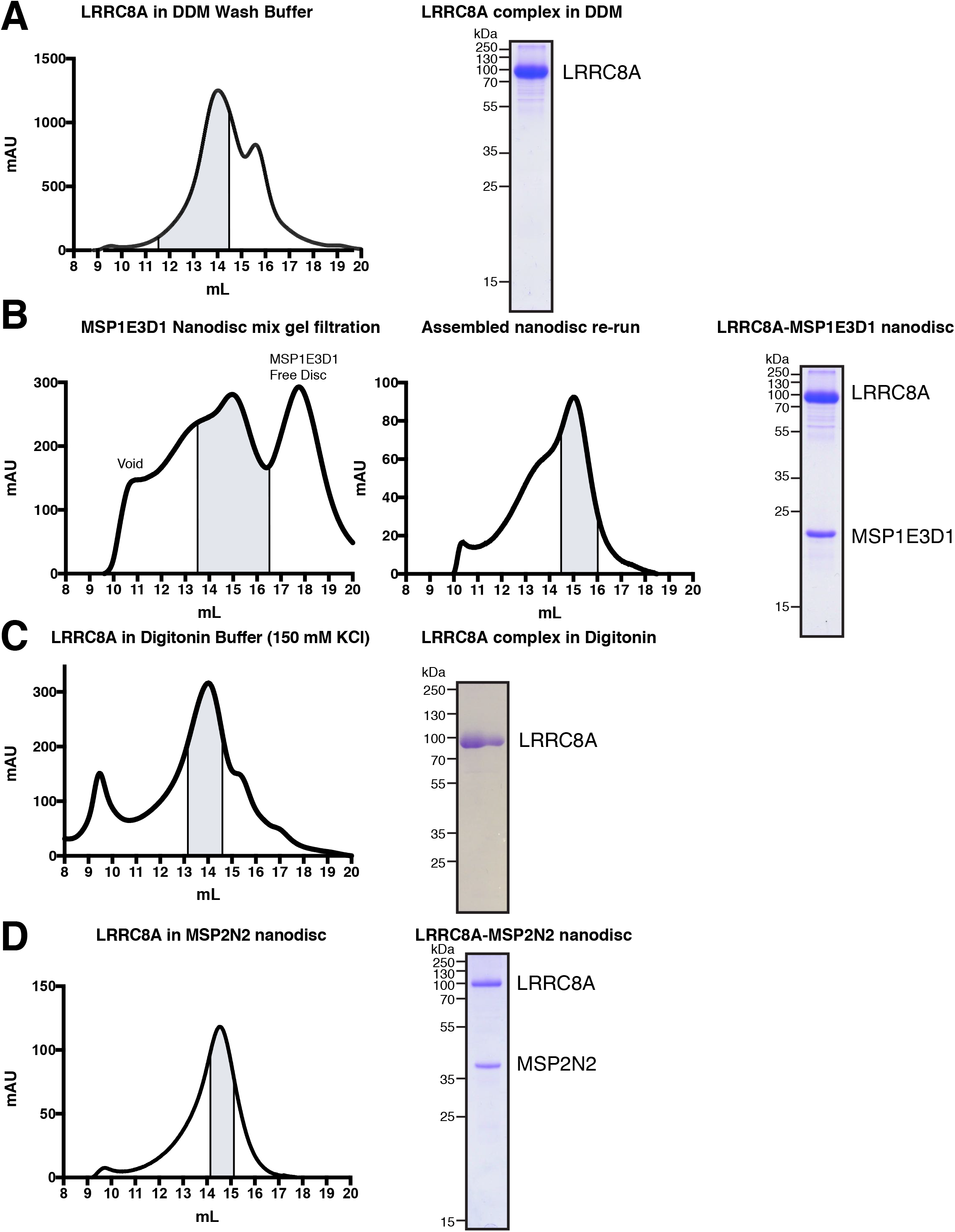
Complex purification and reconstitution. (A) Superose 6 gel filtration of LRRC8A into DDM-containing wash buffer. Peak fractions corresponding to LRRC8A complex are highlighted. On the right, coomassie gel of purified LRRC8A protein in DDM. (B) First run of nanodisc mix on Superose 6 column. Peak fraction corresponding to LRRC8A in MSP1E3D1 nanodiscs is highlighted. These fractions were concentrated and re-run on the same column (middle). Indicated fractions were then pooled and concentrated for grid preparation and analysis by coomassie gel (Right). (C) Superose 6 gel filtration of LRRC8A into digitonin Buffer. Chosen fractions corresponding to LRRC8A complex in digitonin are highlighted. On the right, coomassie gel of purified LRRC8A in digitonin Buffer. (D) Superose 6 gel filtration of LRRC8A in MSP2N2 nanodiscs. Chosen fractions are highlighted. On the right, coomassie gel of purified LRRC8A in MSP2N2 nanodiscs.

**Figure 1-Supplement 2.**
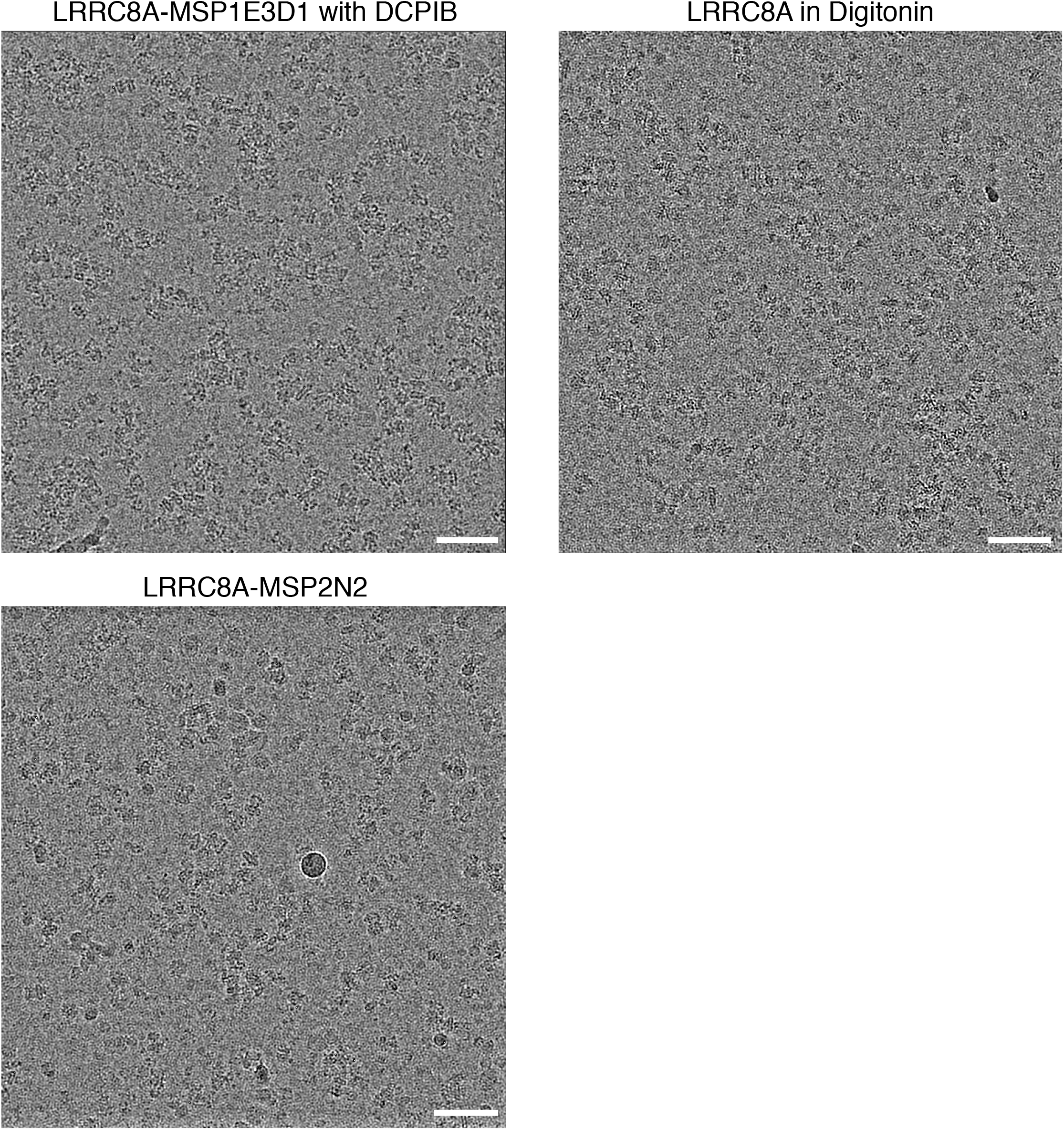
Representative micrographs from LRRC8A preparations in MSP1E3D1 nanodiscs, digitonin, and MSP2N2 nanodiscs. Scale bars, 500 Å.

**Figure 1-Supplement 3.**
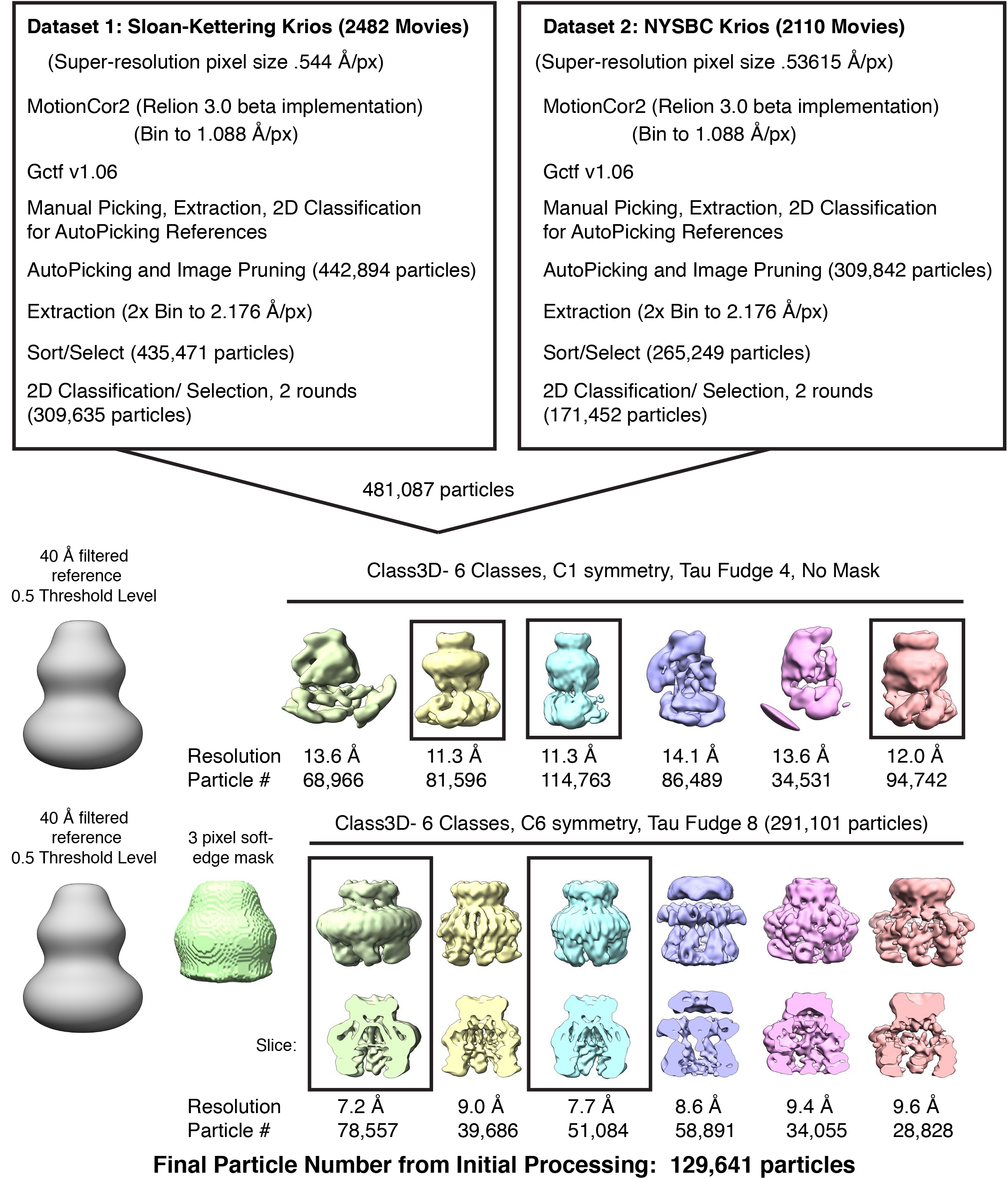
Initial processing for LRRC8A-DCPIB in MSP1E3D1 nanodisc datasets. Particles from boxed classes were passed to the next round.

**Figure 1-Supplement 4.**
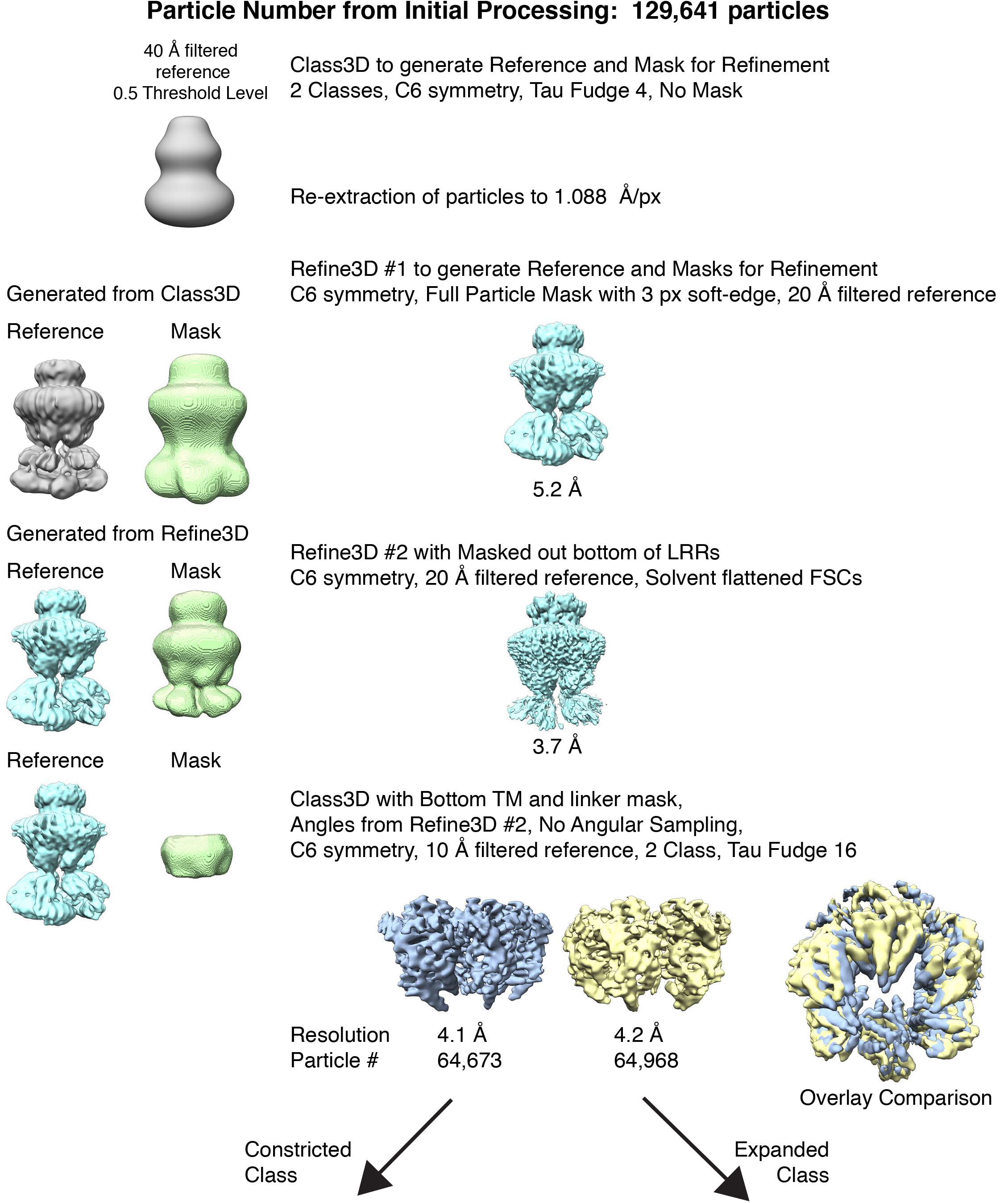
The refinement and classification performed to separate constricted and expanded particles.

**Figure 1-Supplement 5.**
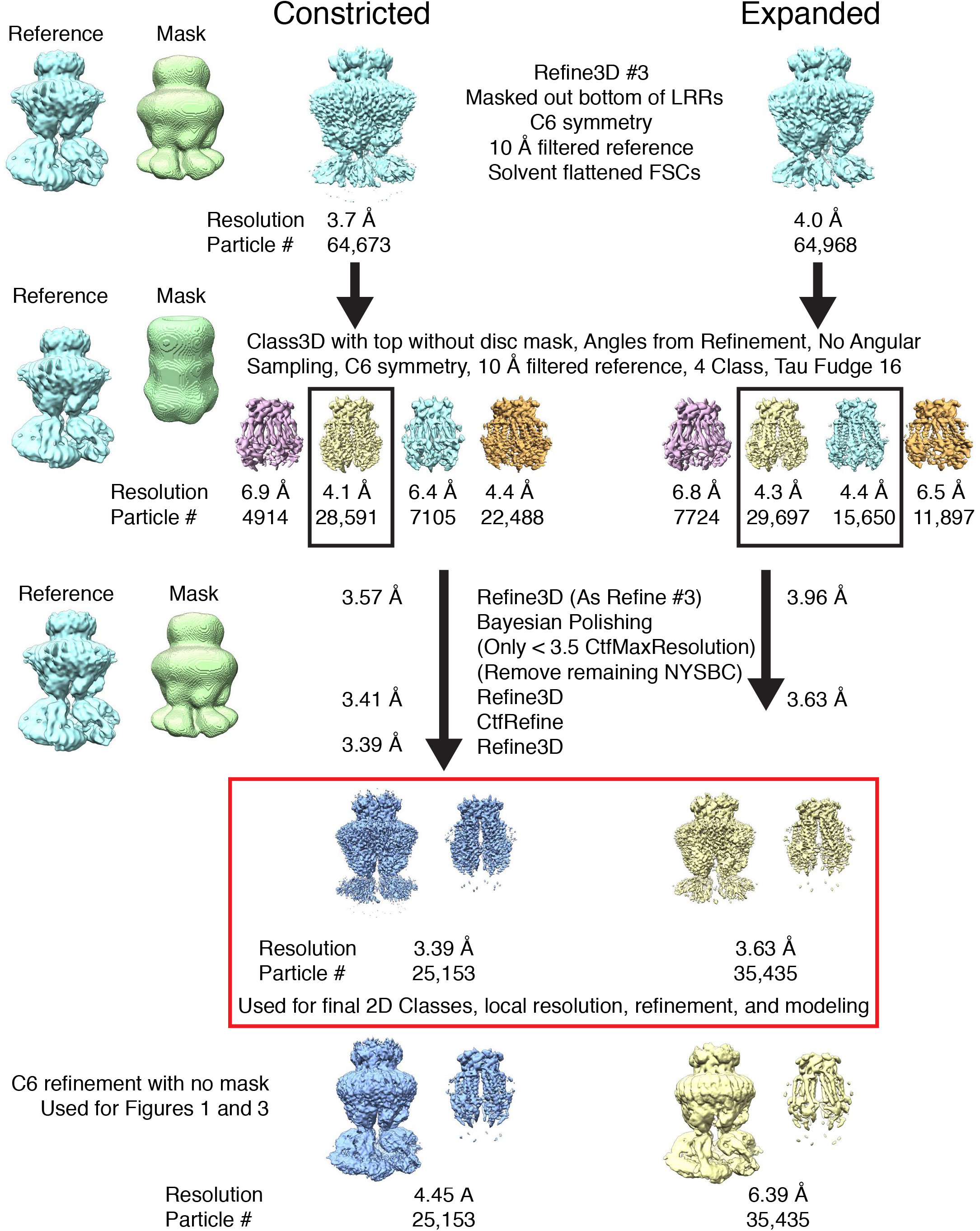
The final classification and refinement performed on constricted and expanded particles to obtain high-resolution maps. Particles from boxed classes were passed to the next round. The masked refinements used for generating final maps and modeling are boxed in red. Below are the unmasked refinements used in Figure 1 and 4.

**Figure 1-Supplement 6.**
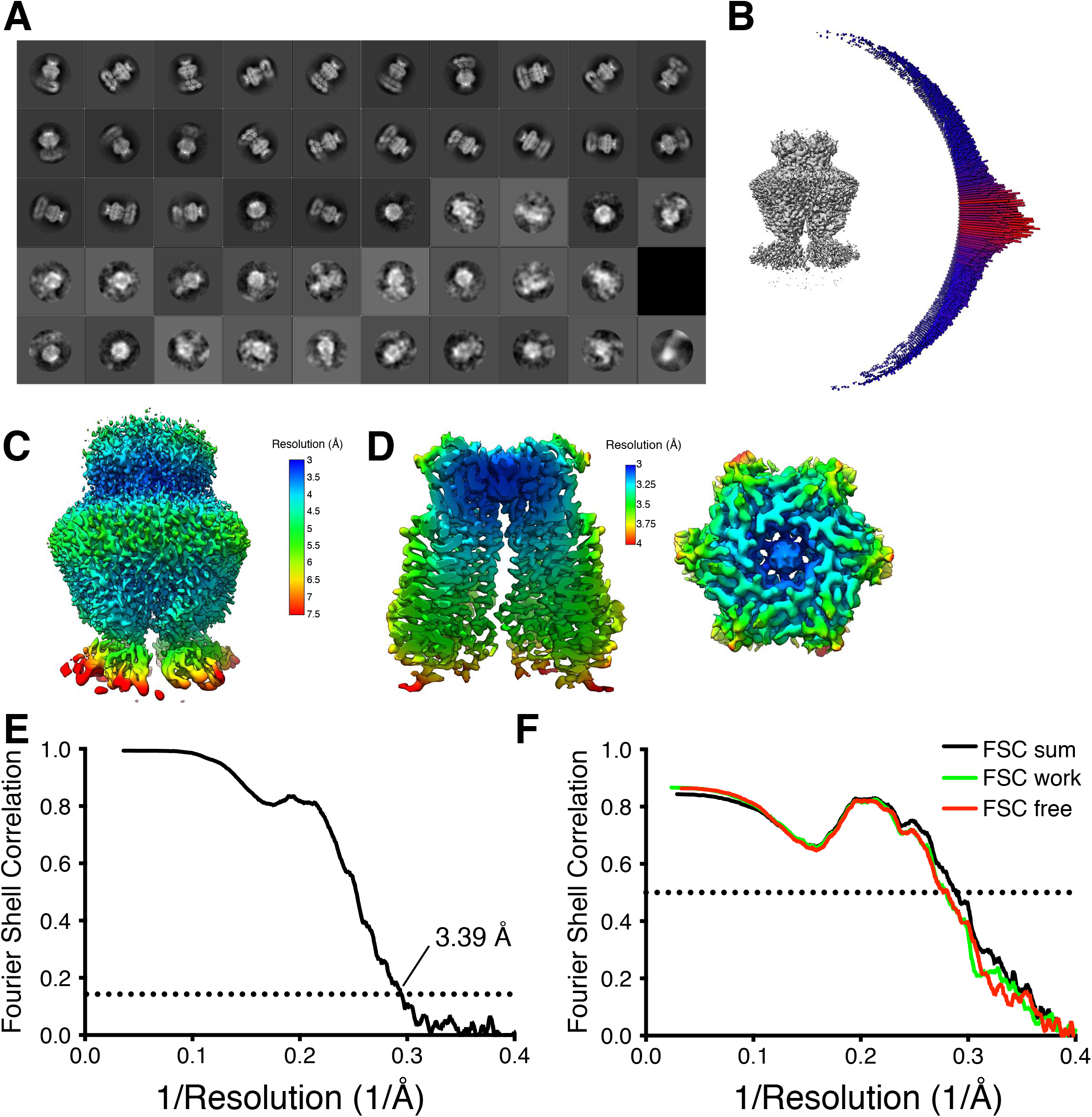
Constricted map and model validation. (A) Two-dimensional classes from a fifty-class classification of the final particles. (B) Angular distribution of particle views. (C) Local resolution using Relion locally filtered map at 0.012 threshold showing resolution of the top of LRR region. (D) Local resolution using Relion locally filtered map at 0.045 threshold and dust hidden at 10 Å. Membrane view (left) and extracellular view (right). (E) Fourier Shell Correlation (FSC) of the two unfiltered half-maps from refinement used for calculating overall resolution at 0.143. (F) Model validation FSC using half-map 1 used for model refinement (FSC work), half-map 2 not used for refinement (FSC free), and full map (FSC sum).

**Figure 1-Supplement 7.**
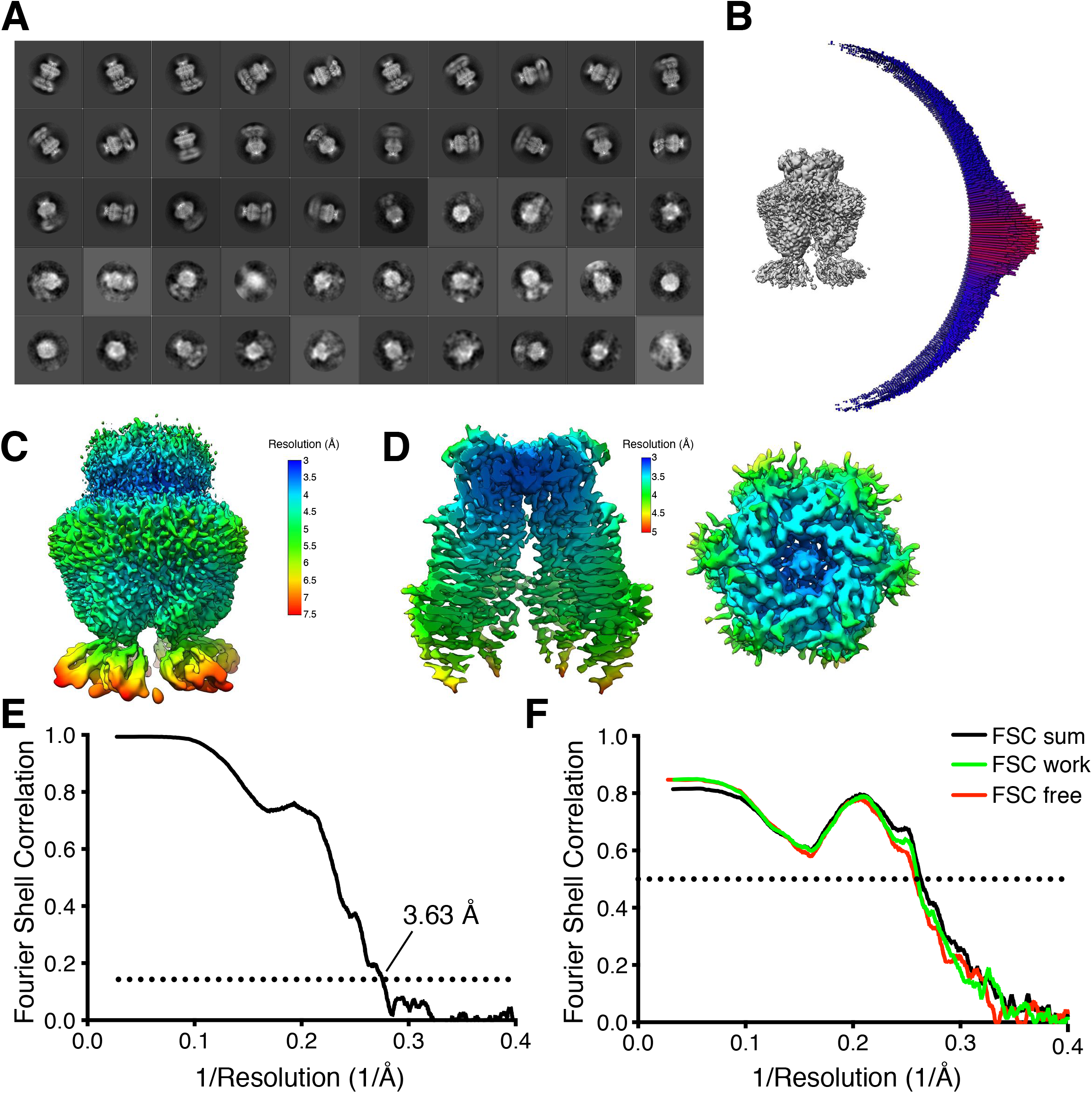
Expanded map and model validation. (A) Two-dimensional classes from a fifty-class classification of the final particles. (B) Angular distribution of particle views. (C) Local resolution using Relion locally filtered map at 0.012 threshold showing resolution of the top of LRR region. (D) Local resolution using Relion locally filtered map at 0.035 threshold and dust hidden at 12 Å. Membrane view (left) and extracellular view (right). (E) Fourier Shell Correlation (FSC) of the two unfiltered half-maps from refinement used for calculating overall resolution at 0.143. (F) Model validation FSC using half-map 1 used for model refinement (FSC work), half-map 2 not used for refinement (FSC free), and full map (FSC sum).

**Figure 4-Supplement 1.**
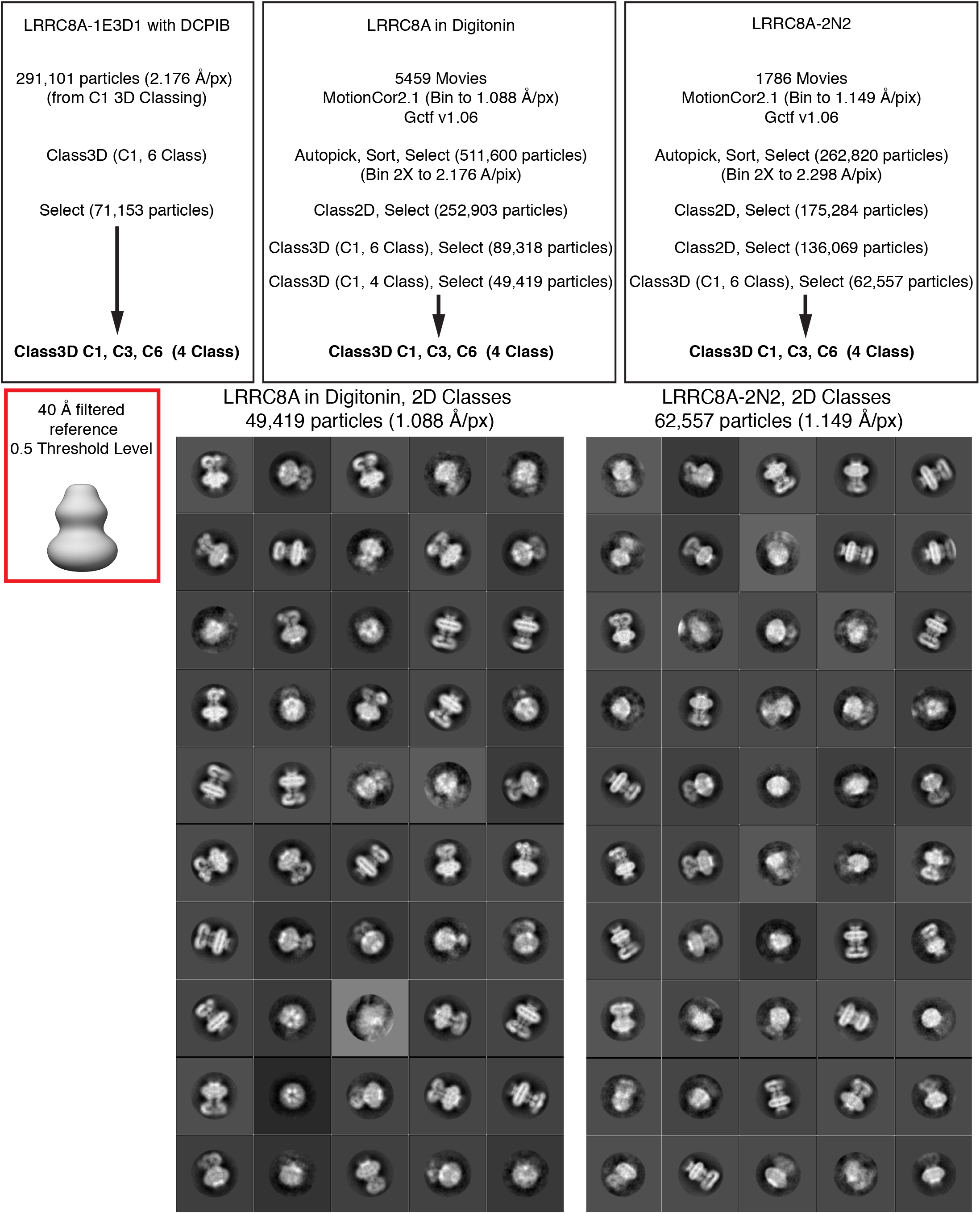
Initial processing for symmetry testing. (Top) The reference used for the three-dimensional classification is shown boxed in red. No masking was utilized during classification. (Bottom) Two-dimensional class averages from a fifty-class job for the final particle set are shown for digitonin and MSP2N2 date.

**Figure 4-Supplement 2.**
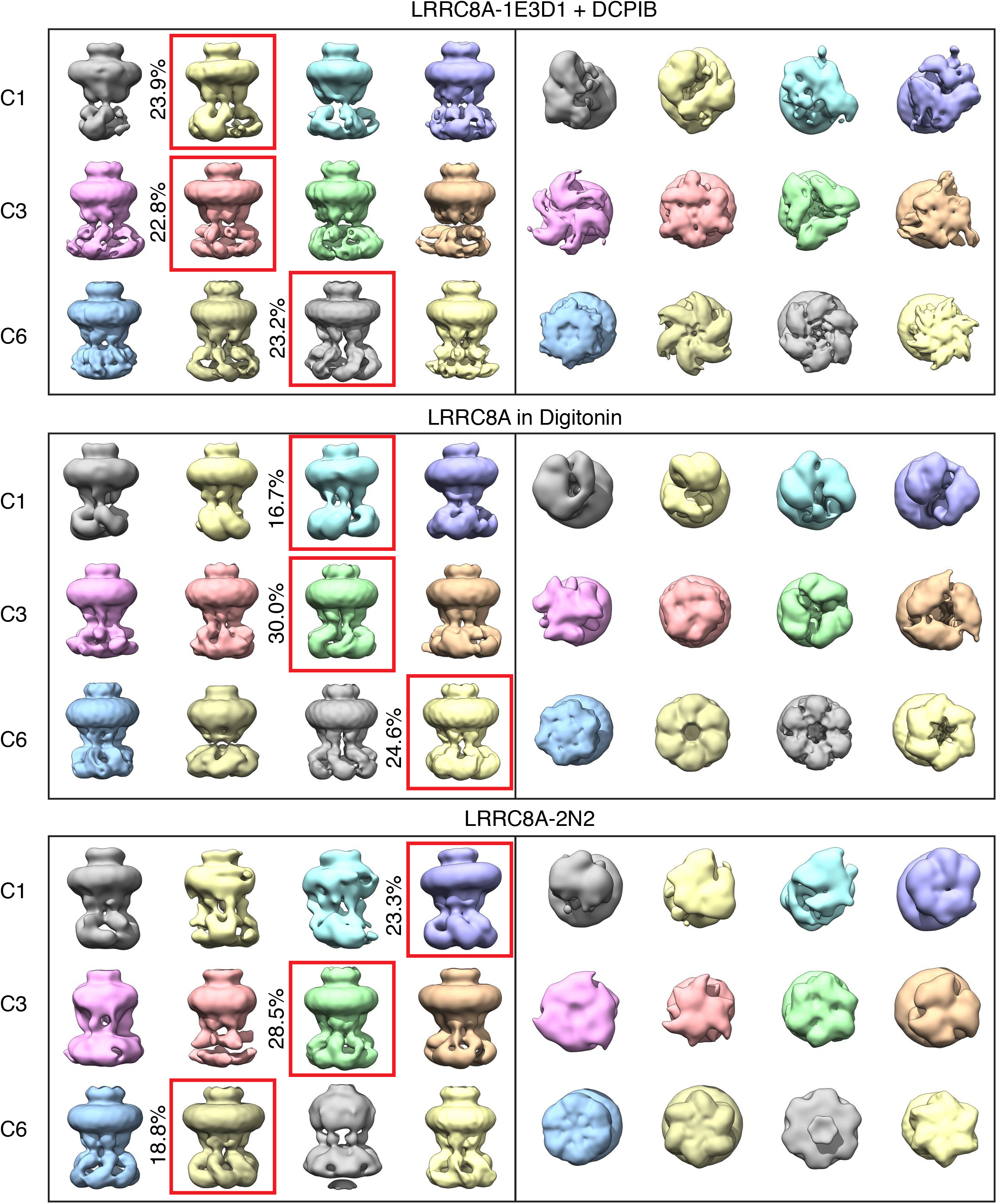
Full three-dimensional classification output for symmetry testing. All maps are shown at .015 threshold. Classes selected for display in Figure 4 are boxed in red with the percentage of particles contributing to the class on the left.

**Movie 1. Motion of two opposing chains (as in Figure 3A) as a cartoon-representation morph between constricted and expanded states.** Measurement in Å between the C-alpha of Pro15 is included.

**Movie 2. Linker region (as in Figure 3C) motion as a cartoon-representation morph between constricted and expanded states.**

**Movie 3. Constricted state side-views illustrating heterogenous LRR positions Movie 4. Expanded state side-views illustrating heterogenous LRR positions**

**Movie 5. Overlay of the constricted state and narrow and wide interfaces of PDB: 5zsu**

**Movie 6. Overlay of the constricted state and narrow and wide interfaces of PDB: 6djb**

